# Epigenomic regulation of human oligodendrocyte myelination properties – relation to age and lineage

**DOI:** 10.1101/2025.09.04.674347

**Authors:** Abdulshakour Mohammadnia, David Dansu, Qiao-Ling Cui, Moein Yaqubi, Roy Dudley, Jo Anne Stratton, Stephanie Zandee, Timothy E. Kennedy, Myriam Srour, Patrizia Casaccia, Jack P. Antel

**Author notes:** Jack P Antel, **Email:**.

## Abstract

Multiple sclerosis (MS) is characterized by immune-mediated injury to myelin and oligodendrocytes (OLs). Repair depends on the ability of OL lineage cells to form new myelin and ensheathe axons. We previously showed that late progenitors (O4+A2B5+ cells) and mature human OLs exhibit age-related differences in ensheathment capacity and vulnerability to injury. Here, we test the hypothesis that differences in chromatin accessibility and specific histone marks may underlie transcriptional differences linked to these functional responses. Confocal imaging of cultured cells revealed higher levels of the transcriptionally permissive histone marks H3K27ac and H4K8ac in pediatric than adult derived late progenitors and mature OLs. Levels were higher in adult-derived progenitors versus mature cells from the same individuals. The majority of pathways and genes related to myelination and immune interactions were downregulated in adult cell samples when compared to pediatric samples. Analysis of publicly available datasets indicated that the chromatin accessibility for genes within these categories was more restricted in adult than pediatric OLs. There was less chromatin accessibility and lower H3K27ac chromatin occupancy also in more differentiated OL compared to progenitors. The levels of the transcriptionally repressive H3K27me3 histone mark in mature OLs were enriched in genomic regions encoding for transcriptional inhibitors of myelination and related signaling pathways, as compared to early progenitors. Restrictions in chromatin accessibility were more pronounced in human cells than in mouse cells. These results link the myelination capacity and immune-mediated injury susceptibility of human OLs to their epigenomic state, raising the issue of how epigenetic modulation could influence disease progression.

**Significance Statement:** Neurological disability in multiple sclerosis reflects a balance between the extent of tissue injury and repair. Human oligodendrocytes display distinct donor age- and maturity stage-related epigenomic profiles, which may influence both their myelination potential and vulnerability to immune-mediated injury. Our results emphasize that epigenomic status, particularly chromatin accessibility and specific histone modifications (H3K27ac and H3K27me3), may underlie these functional capacities. Thus, therapeutic strategies aimed at epigenomic modulation must be carefully considered, due to their net effect on these competing processes—promoting myelin repair while potentially altering susceptibility to further damage—to achieve beneficial clinical outcomes.

## Introduction

Oligodendrocyte (OL) lineage cells ranging from early progenitor cells (OPCs) to mature cells with myelin processes are present within the parenchyma of the developed human CNS. The axon-ensheathing myelin processes not only are a requisite for saltatory conduction but also contain channels permitting transport of essential nutrients to the axons (1). Multiple sclerosis is the prototype acquired human CNS disease in which the OL/myelin unit is considered the primary target of immune-mediated injury. Overall neurologic disability likely reflects the balance of extent of injury and extent of subsequent tissue repair. In MS, extent of myelin repair is variable being more efficient in early disease stage, in younger individuals, and in grey versus white matter (2–7). Although most experimental models with toxin induced destruction of mature OLs indicate the role of OPCs in underlying subsequent remyelination (8), other models indicate that mature OLs can be responsible (9–11). Mature OL cell bodies are relatively preserved in early MS lesions (2). Xenogeneic transplant studies indicate that progenitors far along in the OL lineage, O4+ cells recognized by A2B5 antibody and referred to as “pre-OLs can myelinate the genetically dysmyelinated CNS of shiverer mice; the extent of myelination per cell was found to exceed that mediated by fetal early OPCs (12–14). As with myelination capacity, OL interaction with immune system is also lineage stage dependent. Lineage progenitors express an increase in immune related molecules that contribute to their enhanced vulnerability to immune injury compared to mature OLs (15, 16).

In our previous studies using A2B5+O4+ cells (late progenitors) and mature OLs derived from surgical resection of human brain, we observed that ensheathment of synthetic nanofibers was greater for cells from pediatric versus adult donors consistent with in situ studies cited above (17). For both age groups, we found that ensheathment was greater for the late progenitors compared to mature OLs. Bioinformatics analyses of these data sets showed that most variance between the cell types measured on day one after isolation was due to age (43% of variance on PC plot) with secondary contribution from lineage (17%). We further observed that our cell populations with superior ensheathment also showed enhanced susceptibility to immune and stress conditions mediated injury (18).

The equilibrium between histone acetylation and deacetylation provides a major means of epigenomic modulation of oligodendrocyte differentiation (19, 20). Histone acetylation is associated with increased levels of transcriptional inhibitors of oligodendrocyte differentiation whereas deacetylation favors differentiation (21). Shen et al (22) showed that in demyelinated young mouse brains, new myelin synthesis was preceded by downregulation of oligodendrocyte differentiation inhibitors and neural stem cell markers, and that this was associated with recruitment of histone deacetylases (HDACs) to promoter regions. In demyelinated old brains, HDAC recruitment was inefficient, and this allowed the accumulation of transcriptional inhibitors and prevented the subsequent surge in myelin gene expression. Pedre et al., (2011) reported a shift toward histone acetylation, in the white matter of the frontal lobes of aged subjects and in patients with chronic multiple sclerosis (23). The data in chronic lesions contrasted with findings in early MS lesions, where marked oligodendroglial histone deacetylation was observed (23). Liu et al found that that oligodendrocytes in tissue sections containing active and inactive MS lesions are epigenetically silenced (low or undetectable levels of H3K27ac) and lack myelin production (24). They further showed that inhibiting HDAC3 with ESI1 leads to strong ectopic myelination in a mouse model, correlating with higher levels of H3K27ac enhanced in vitro ensheathment of nanofibers by rodent pre-OLs.

To address the postulate that age and lineage maturity linked changes in myelination and immune properties in human OL lineage cells reflect epigenomic status, we first used immunostaining of primary human mature and late lineage progenitor cells (O4+A2B5+) to document that levels of the transcriptionally permissive H3K27ac histone modification were higher in pediatric than adult samples, in both late progenitors and mature OLs. We then used our RNA dataset of pediatric and adult samples of late progenitors and mature OLs to derive an age-related molecular signature relevant for myelination, immune, and stress responses for each of these cell types and further determined whether these signatures also linked with lineage stage. We further used publicly available data sets to show that this signature is regulated at the epigenomic level as measured by levels of chromatin accessibility and the active enhancer histone mark H3K27ac both in relation to age and lineage stage. In view of the significant available studies on properties of murine derived OL lineage cells we compared our human based observations with biologic properties of rodent cells. Our results link both myelination capacity and immune-injury susceptibility of human OLs to epigenome state, raising a word of caution regarding the net effect of epigenetic modulation as potential therapeutic venue.

## Results

### Age linked changes in transcriptome and epigenome of human OLs

**Immunofluorescence based analysis of transcriptionally permissive (H3K27ac and H4K8ac) and repressive (H3K27me3) histone marks in human late progenitors and mature OLs.**

To asses age-related epigenome changes linked to histone-post-transcriptional modifications, we investigated the levels of marks linked to transcriptional activation, such as the active enhancer mark H3K27ac, as well as H4K8ac, which have been shown to decrease with age in mice (25), and marks linked to heterochromatin and transcriptional silencing, such as H3K27me3. Immunofluorescence staining, followed by quantitative confocal analysis demonstrated an age-dependent global reduction of all the assessed marks, including H3K27ac (Fig. 1A-D, Fig. S1A), H4K8ac (Fig. S1C, Fig. S2A-D) and H3K27me3 (Fig. 1E-H, Fig. S1B) levels in both late progenitor cells and in mature OLs during the pediatric-to-adult transition (Fig. 1). This suggests that the ability of human cells to modulate epigenomic marks decreases overall with age.

**Figure 1.**
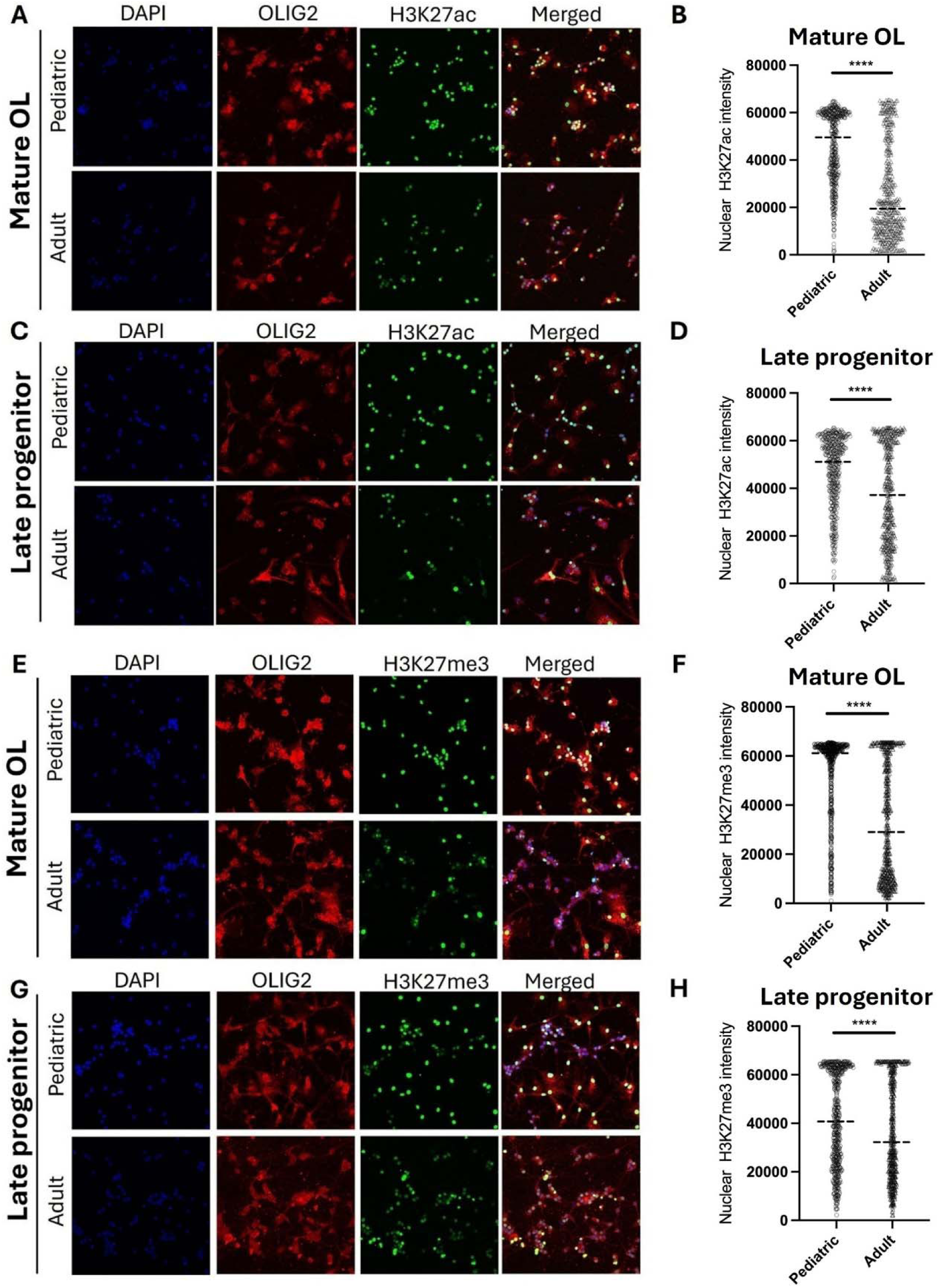
H3K27ac and H3K27me3 profiling in late human progenitor A2B5+ and mature A2B5− cell in both pediatric and adult groups. **(A)** Confocal image of A2B5^−^ cells obtained from pediatric and adult human patients, stained with OLIG2 antibodies (red), H3K27ac antibodies (green) and DAPI (blue) as a nuclear counterstain. **(B)** Violin plots of the nuclear intensity of H3K27ac immunoreactivity in OLIG2^+^ cells in pediatric and adult brains. Data represent the H3K27ac immunoreactivity measured in a total of 450 cells from three biological replicates (****P < 0.0001, unpaired t-test).. **(C)** Confocal image of A2B5^+^ cells obtained from pediatric and adult human patients, stained with OLIG2 antibodies (red), H3K27ac antibodies (green) and DAPI (blue) as a nuclear counterstain. **(D)** Violin plots of the nuclear intensity of H3K27ac immunoreactivity in OLIG2^+^ cells in pediatric and adult brains. Data represent the H3K27ac immunoreactivity measured in a total of 450 cells from three biological replicates (****P < 0.0001, unpaired t-test). **(E)** Confocal image of A2B5^−^ cells obtained from pediatric and adult human patients, stained with OLIG2 antibodies (red), H3K27me3 antibodies (green) and DAPI (blue) as a nuclear counterstain. **(F)** Violin plots of the nuclear intensity of H3K27me3 immunoreactivity in OLIG2^+^ cells in pediatric and adult brains. Data represent the H3K27me3 immunoreactivity measured in a total of 600 cells from three biological replicates (****P < 0.0001, unpaired t-test). **(G)** Confocal image of A2B5^+^ cells obtained from pediatric and adult human patients, stained with OLIG2 antibodies (red), H3K27me3 antibodies (green) and DAPI (blue) as a nuclear counterstain. **(H)** Violin plots of the nuclear intensity of H3K27me3 immunoreactivity in OLIG2^+^ cells in pediatric and adult brains. Data represent the H3K27me3 immunoreactivity measured in a total of 600 cells from three biological replicates (****P < 0.0001, unpaired t-test).

### Age linked changes in transcriptome and epigenome of human OLs

To link function with epigenome, we first generated a new overall comparative age-related gene expression signature using our previously published age-related bulk RNA-seq data (17) derived from late progenitor cells (A2B5+) and mature (A2B5−) OLs one day after isolation, the earliest time point at which we could derive a pure population suitable for bulk RNA sequencing (Fig. 2A). Overall, there were 986 protein coding genes commonly downregulated in OL lineage cells (both late progenitors and mature cells) in adult samples compared to pediatric samples (Fig. 2B). 1151 and 263 were specifically downregulated in late progenitors and mature cells (p < 0.05 and Log2FC >1) by age respectively (Fig. 2B, Fig. S3). Gene ontology (GO) enrichment analysis of the down-regulated coding genes common to both cell types performed using Gprofiler, with subsequent visualization in Revigo (Fig. 2C, Fig. S3A-B), identified significantly enriched biological processes (adjusted p<0.05) then categorized into three major functional groups (Fig. 2C, Supplementary File S1, Fig. S1A-B). The “myelination-related signature” (MRS) included terms and genes related to OL differentiation, including genes associated with the progenitor stage, such as *PTPRZ1* and *SOX5* and broadly defined myelination-relevant pathways, including those regulating glial cell differentiation, cell adhesion, and additional signaling pathways (Supplementary file 1); of note, this signature did not include myelin-structural genes such as *MAG*, *MOG*, and *PLP1*. The additional categories included “immune-related signature” (e.g., innate immune response, inflammatory response, response to virus (Supplementary file 1) and “stress response and cell death-related processes” (e.g., response to stress, programmed cell death, apoptosis; supplementary file 1) (Fig. 2C). From these three broad categories, we extracted specific genes reflecting each category: 119 genes from the “myelination-related” signature, 40 genes from the “immune-related” signature and 81 genes from the “stress/cell death related” processes (Fig. 2D, Supplementary file 1) and asked whether we could detect significant difference between pediatric and adult samples. Single-sample gene set enrichment analysis (ssGSEA) revealed a significant enrichment (*p<0.05*) in pediatric samples, across both late progenitors and mature OLs (Fig. 2E). We showed the same trends for our signatures using an independent dataset in which cells were kept for six days in culture (26) (Fig. S4A). In comparison with downregulated genes, only 253 and 498 genes were specifically upregulated in late progenitors and mature cells by age respectively (Fig. S3C, Fig. S3E-F). There was no specific enrichment for the relevant pathways of the 3 distinct categories considered in the previous section.

**Figure 2.**
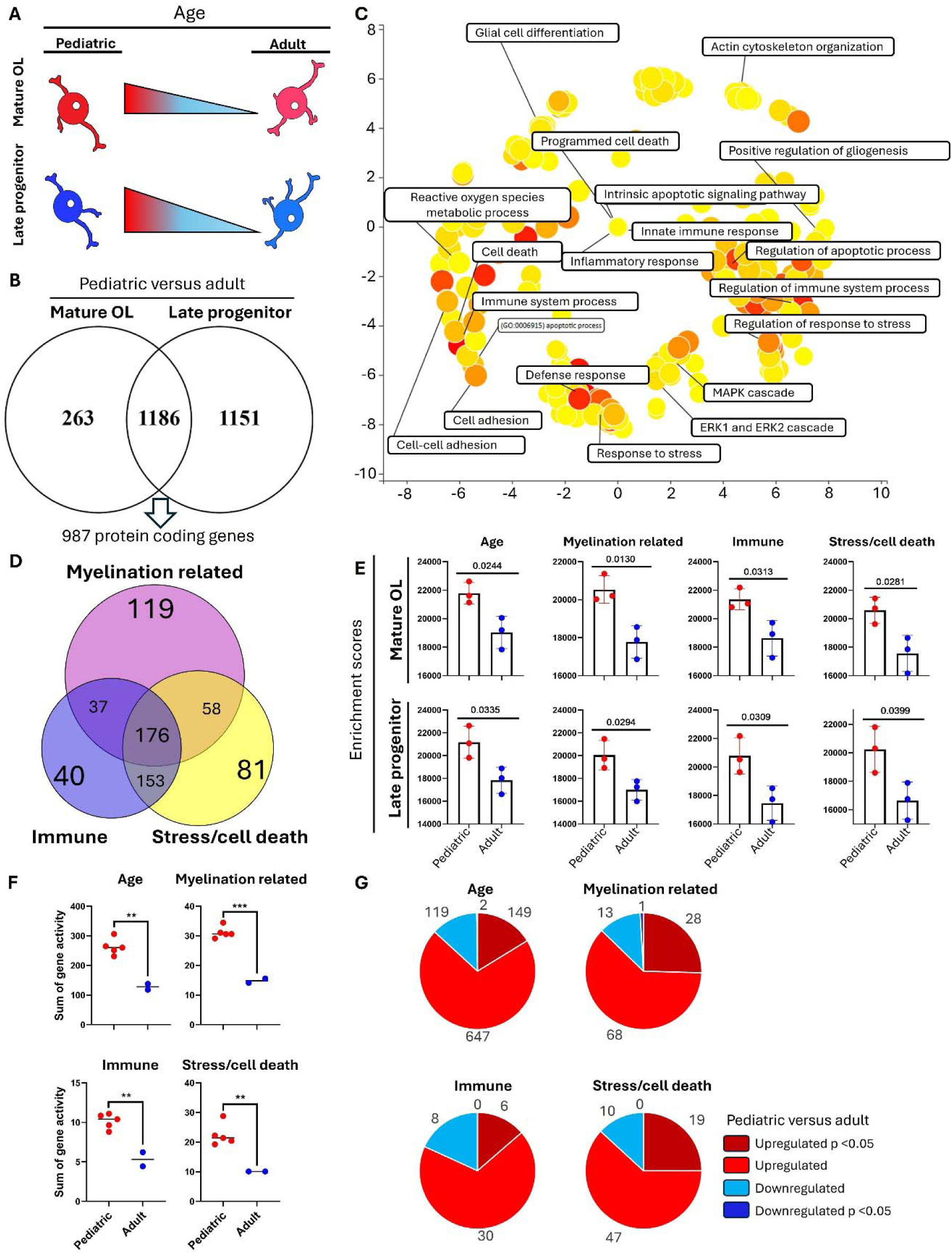
Transcriptomic signature is downregulated in mature oligodendrocytes and late progenitors in adult samples compared to pediatric samples. **(A)** Age-related transcriptomic signatures were identified by comparing adult (>18 years) and pediatric (<18 years) samples in two distinct human oligodendrocyte lineage populations: mature oligodendrocytes (A2B5−) and late oligodendrocyte progenitors (A2B5+). Triangles reflect ensheathment potential which is higher in pediatric samples for both late progenitors and mature cells in comparison to their adult counterparts. **(B)** Genes downregulated in adult samples for each comparison were filtered based on statistical significance (p < 0.05) and log2 fold-change (>1), resulting in a set of differentially expressed genes. Intersection of these sets yielded 1186 overlapping genes; removal of non-coding genes refined this list to 987 protein-coding genes. **(C)** Gene ontology (GO) analysis was performed using Gprofiler on the 987 genes, and significant GO terms (adjusted p < 0.05) were visualized using a scatter plot generated in Revigo, with representative significant terms highlighted. **(D)** Significant GO terms were classified into three categories: (1) Myelination related signature (MRS) which includes oligodendrocyte lineage differentiation and myelination related processes), (2) immune-related processes, and (3) stress and cell death. Genes specific to each category were identified for further analysis. **(E)** Single-sample gene set enrichment analysis (ssGSEA) was conducted on mature oligodendrocyte and late progenitor expression profiles for the identified gene categories (full age-related signature, myelination related, immune, and stress/cell death signatures). Enrichment scores were plotted, and statistical significance between pediatric and adult groups was assessed using t-tests. All signatures showed significantly higher enrichment scores in pediatric samples compared to adults. **(F)** The ATAC-seq modality of the single-cell multiomic dataset from Velmeshev et al. (2023), including pediatric (ages 1–8 years) and adult (25 and 39 years old) donors, was used to assess accessibility (gene activity) of gene signatures derived from comparing bulk RNA-seq adult human primary cells with pediatric counterparts. For each sample, average gene activity for genes in each group for mature OLs was calculated, and the sum of gene activities per signature per sample was used. A two-sided t-test was used to assess the significance of differences between adults and pediatrics; ** and *** indicate p < 0.005 and p < 0.0005, respectively. **(G)** We performed a pseudobulk differential analysis on mature OLs (comparing pediatric versus adult) by aggregating normalized gene-activity values (log(CP10k)) to donor-level means, using Individual as the biological replicate. Donors were split into pediatric (age 1 to 8 years old) and adult groups (age > 20), and per-gene differences were tested with Welch’s t-test to accommodate unequal variances and sample sizes. Genes from the overall age and 3 sub-category signatures were extracted and divided into four categories: significantly upregulated (p < 0.05, dark red), upregulated (non-significant, light red), downregulated (non-significant, light blue), and significantly downregulated (p < 0.05, dark blue).

To correlate our age related signature gene expression with age dependent chromatin accessibility, we used the ATAC seq modality of a single cell multiomic postnatal dataset provided in Velmeshev et al. (2023) (27, 28). Specifically, we used early pediatric samples (from 1 year up to 8 years old, (comparable with our pediatric samples) and adult samples (older than 20 years) (Fig. 2F and 2G). Enrichment analysis revealed a significant age-dependent decrease in chromatin accessibility in OLs for the overall age-related signature as well as for all three subcategories of myelination related, immune, and stress/cell death (Fig. 2E). Quantifying gene activity and calculating differences and significance for genes in our signatures revealed that most genes have higher accessibility (gene activity) in pediatric mature OLs compared with adult counterparts (Fig. 2G). Early oligodendrocyte progenitors (OPCs) from this data set showed the same trend (pediatric versus adult), with lower enrichment for the different signatures by age, and the majority of genes showed lower accessibility in adult OPCs (Fig. S4B and S4C). Overall chromatin accessibility, and specifically for each of our gene categories including the myelination related genes, was more restricted in OLs from adult versus pediatric samples, reflecting that decreasing gene expression with age correlates with decreasing accessibility at the epigenome level.

### Lineage linked changes in transcriptome and epigenome of human OLs

To establish whether our observed age-related myelination related s ()(correlates with higher level of ensheathment Fig. 2A), immune, and stress signatures also related to lineage or maturity stage, we analyzed three publicly available snRNA-seq datasets ((29–32) derived from adult human tissue samples, together with our own scRNA-seq dataset (30). Within these datasets, oligodendrocyte lineage cells, including OPCs and mature oligodendrocytes, were identified and annotated. Average expression levels extracted from OPCs and mature cells were subjected to enrichment analyses. SsGSEA for full age signature and the three subsets (myelination related, immune, and stress/cell death) consistently revealed higher enrichment in OPCs than mature oligodendrocytes (Fig. 3A). Calculated fold changes between OPCs and mature oligodendrocytes based on average expression values confirmed higher expression in OPCs for many genes across signatures (87.8% for total age-related, 90.7% for myelination related, 89.5% for immune, and 85.4% for stress/cell death signatures) (Fig. 3B). Examples of genes showing higher expression both in pediatric versus adult cells and in OPCs compared to mature OL include progenitor markers (e.g. *PTPRZ1, SOX5*), as well as immune related genes (e.g. *DRAM1, TNFRSF1A, and HLA-B*) are provided in Supplementary file 2, Fig. 3C-D.

**Figure 3.**
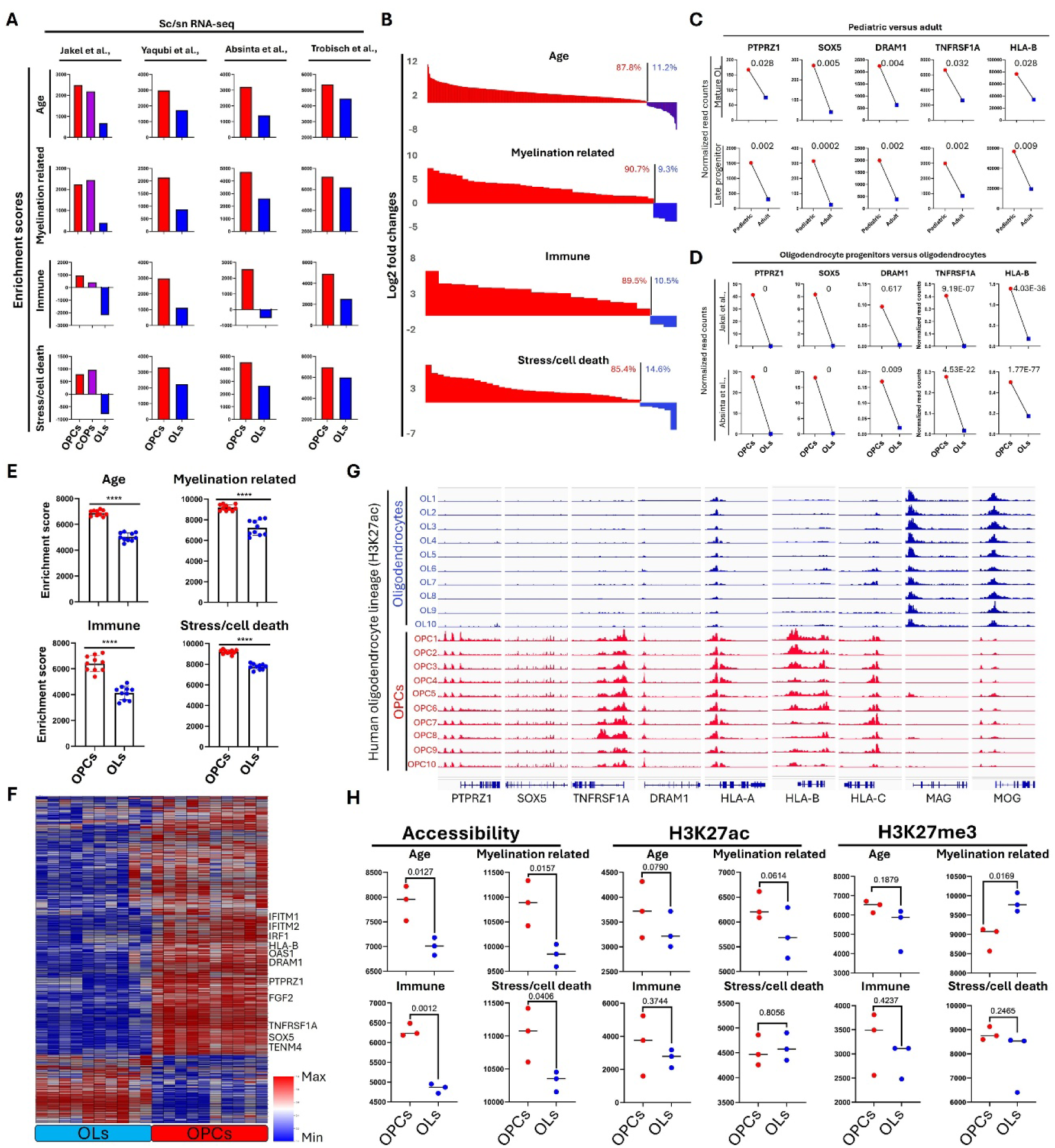
Age-related transcriptomic and epigenomic signatures are upregulated in oligodendrocyte progenitor cells (OPCs) and downregulated in mature oligodendrocytes (OLs). **(A)** Gene expression profiles of OPCs and mature OLs were analyzed using three single-nucleus RNA sequencing (snRNA-seq) datasets (Jakel et al., 2019; Absinta et al., 2021; Trobisch et al., 2022) and one single-cell RNA sequencing (scRNA-seq) dataset (Yaqubit et al., 2023). Jakel et al. (2019) additionally included committed oligodendrocyte progenitors (COPs). Average gene expression from OPCs and mature OLs across these datasets served as input for single-sample gene set enrichment analysis (ssGSEA) for age-related, OL differentiation (correlate with higher nanofiber ensheathment potential), immune, and stress/cell death signatures. Calculated enrichment scores revealed consistently higher signature enrichment in progenitor cells and decreased enrichment in mature cells. **(B)** Gene expression profiles were extracted from the Jakel et al. (2019) dataset for all signatures, and fold changes were plotted. Higher expression in OPCs is indicated by red, whereas higher expression in mature OLs is shown in blue. The percentage of genes upregulated or downregulated within each signature was also plotted. **(C)** Representative genes involved in OL differentiation and myelination (PTPRZ1, SOX5) and in stress, cell death, and immune functions (DRAM1, TNFRSF1A, HLA-A) exhibited significantly higher expression in pediatric samples (both late progenitors and mature cells) compared to adult samples (p<0.05). *P-values* for gene expression were calculated using the DESeq2 package in R. Normalized read counts are depicted, with pediatric samples in red and adult samples in blue. **(D)** The same representative genes also showed significantly higher expression in OPCs compared to mature OLs (p<0.05). Normalized read counts are plotted with OPC expression indicated in red and mature cells in blue. DRAM1 and TNFRSF1A exhibit lower gene expression levels compared to other genes in snRNA-seq datasets. *P-values* were calculated using the FindMarkers function in the Seurat package in R. **(E)** ChIP-seq data from Kozlenkov et al. (2024), assessing H3K27ac (active histone modification) at transcription start sites (10 kb) of genes in OPCs and mature OLs from human post-mortem samples, were analyzed. H3K27ac levels quantified using normalized RPKM values were subjected to single-sample gene set enrichment analysis (ssGSEA) for overall age, myelination related, immune, and stress/cell death signatures. Enrichment scores were plotted per sample, with statistical significance (*p-values*) calculated using t-tests. Red indicates OPCs, and blue indicates mature OLs Higher enrichment in OPCs across signatures aligns with the observed gene expression downregulation in mature cells correlating with decreased H3K27ac levels. **(F)** Hierarchical clustering and heatmap visualization of genes with significant differences in H3K27ac levels (p<0.05, affecting 68% of genes in the full age-related signature) at TSS regions. Clustering utilized RPKM values, demonstrating predominant enrichment in OPCs compared to mature cells. Representative genes related to myelination potential (PTPRZ1, SOX5, TENM4, FGF2) and immune and stress/death functions (IFITM1, IFITM2, IRF1, OAS1, HLA-B, DRAM1, TNFRSF1A) are highlighted. Heatmap color scale is row-normalized; yellow indicates higher H3K27ac levels, blue indicates lower levels. **(G)** Integrated Genome Viewer (IGV) tracks illustrating differential H3K27ac enrichment in OPCs (red) versus mature OLs (blue) at representative loci, including myelination related genes (PTPRZ1, SOX5), immune/stress/death genes (TNFRSF1A, DRAM1, HLA-A), and mature OL marker MAG. Tracks were group auto-scaled for visualization. **(H)** Single-cell ATAC-seq and nanoCUT&Tag data for H3K27ac (activating mark) and H3K27me3 (repressive mark) from Kabbe et al. (2024) were analyzed at gene TSS regions (10 kb). ssGSEA performed on RPKM values for accessibility, H3K27ac, and H3K27me3 signatures identified significant differences between OPCs (red) and mature cells (blue). Enrichment scores and statistical significance (t-test) were plotted and annotated.

We then utilized bulk ChIP-seq data to assess H3K27ac on human OPCs and mature cells in the data set from Kozlenkov et al. (2024) (33) H3K27ac levels were quantified within ±10 kb of transcription start sites (TSS), normalized as RPKM values, and subjected to enrichment analysis. This analysis showed consistently higher H3K27ac enrichment for all signatures in OPCs relative to mature oligodendrocytes (Fig. 3E) aligned with the observed higher gene expression levels in OPCs (Fig. 3A). Of all genes within the complete signature, 68% exhibited significant differences in H3K27ac levels around the TSS (p<0.05), with the majority (79%) showing higher levels in OPCs (Fig. 3F). Integrated Genome Viewer (IGV) shows increased H3K27ac levels at genes in OPCs compared to mature oligodendrocytes, including genes related to myelination (*PTPRZ1, SOX5*) and immune/stress/cell death functions (*TNFRSF1A, DRAM1, HLA-A, HLA-B, HLA-C*) (Fig. 3G). Conversely, *MAG* and *MOG*, markers highly expressed in mature oligodendrocytes, display an opposite pattern (Fig. 3G). To further elucidate epigenomic regulation, single-cell resolution analyses of chromatin accessibility (snATAC-seq) and histone modifications (nanoCUT&Tag for H3K27ac as an activation mark and H3K27me3 as a repressive mark) were performed (34). Quantification within ±10 kb TSS regions again demonstrated higher accessibility correlating with higher gene expression in OPCs. H3K27ac levels were overall higher in OPCs and specifically for genes within the “myelination-related” signature, corresponding to progenitor genes and signaling pathways (e.g. *PTPRZ1* and *SOX5)*. Conversely, H3K27me3 showed increased enrichment for the same genes in mature OLs, consistent with the decreased expression of these markers in differentiated cells (Fig. 3H).

### Epigenomic and transcriptomic differences between human and rodent oligodendrocyte lineage cells

Epigenomic differences could underline transcriptomic and functional differences in myelination and response to injury between human cells and rodent cells. As regards transcriptomic features, analysis of bulk RNA-seq data from Neumann et al. (2019) (35) using young (2–3 months) and aged (20–24 months) rat A2B5+ cells indicated that transcriptomic differences between young and aged rodents were minimal for most signatures, with only the myelination related signature showing significant enrichment in A2B5+ cells, suggesting greater similarity between young and aged rodent OL lineage cells compared to their human equivalents (Fig. S5A). Comparing OPC and mature OL marker genes between young rodents versus our human pediatric samples and aged rodents versus our adult human samples demonstrated higher expression of mature markers and lower expression of progenitor markers in human cells (Fig. 4A).

**Figure 4.**
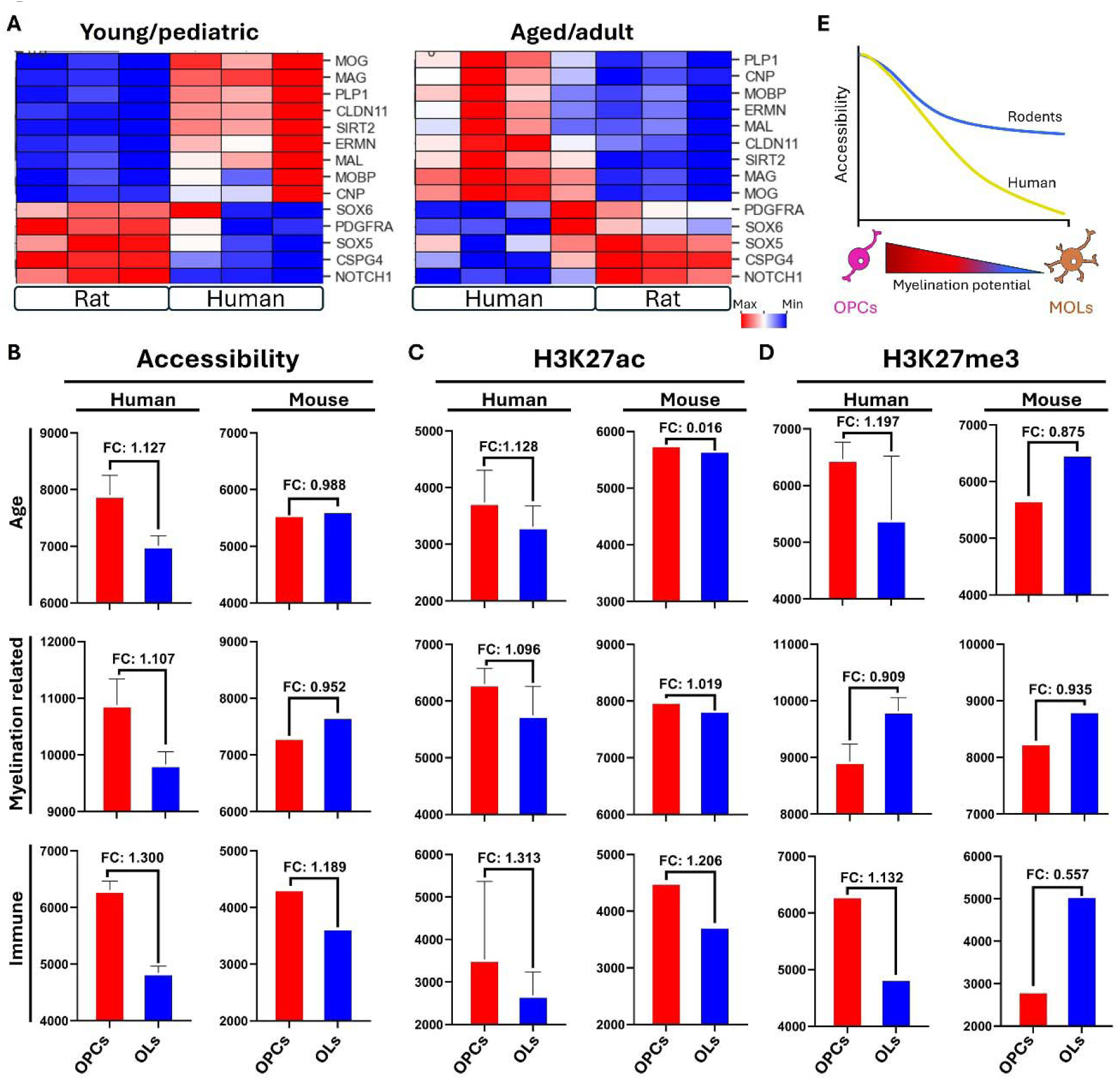
Comparative epigenomic profiling of age-related signatures between human and rodent oligodendrocytes. **(A-C)** Single-cell ATAC-seq and single-cell nanoCUT&Tag datasets for human (Kabbe et al., 2024) and single-cell ATAC-seq and snCUT&Tag datasets for mouse (Bartosovic et al., 2021) were used. Accessibility, H3K27ac (activation mark, n=2, samples pooled), and H3K27me3 (repressive mark, n=4, samples pooled) were quantified within ±10 kb TSS regions across all genes (normalized as RPKM values) in OPCs and mature oligodendrocytes from both species. ssGSEA was performed for three gene signatures (age-related, myelination, and immune). Enrichment scores are plotted with fold differences plotted for the conditions. Red represents OPCs and blue indicates mature cells. **(D)** Transcriptomic comparison between human pediatric (A2B5+) cells and young rat (A2B5+) cells, and human adult (A2B5+) cells and aged rat (A2B5+) cells using normalized read counts. Heatmap visualizations, highlighting markers of OPC and mature oligodendrocytes, indicate higher expression of mature markers in human A2B5+ cells and lower expression of OPC marker genes compared to rat cells. Yellow and blue colors indicate upregulation and downregulation, respectively. Rat transcriptomic data were sourced from Neumann et al. (2019). **(E)** Schematic illustrating the epigenomic status in OPCs and mature oligodendrocytes in humans versus rodents, emphasizing reduced chromatin accessibility for age-related and myelin-related genes in human cells.

For epigenomic analyses, we used single-cell resolution data for chromatin accessibility (ATAC-seq), and histone marks H3K27ac (activation) and H3K27me3 (repression) to compare the epigenome related to our gene signatures between equivalent cell types from humans and rodents; the latter using mouse datasets from Bartosovic et al. (2021) (36). OPCs and mature OL lineage cells were examined in both species, and normalized RPKM values for chromatin features within ±10 kb TSS regions were analyzed using ssGSEA. Chromatin accessibility and H3K27ac levels were higher in OPCs compared to mature cells in both human and rodent datasets, although the magnitude of these differences did not always reach statistical significance in mice (Fig. 4B-D, Fig. S5B). H3K27me3, the repressive histone mark, showed differences in human and rodent cells. While greater levels of this mark were detected in rodent cells for all the transcriptional signatures analyzed, similarly increased H3K27me3 levels were detected in human cells only for “myelination-related” gene signature, during the maturation of OPC into OLs (Fig. 4D, Fig. S5B). This suggests that mechanisms of progenitor gene repression are conserved among species, while intrinsic differences in chromatin accessibility for other transcriptional signatures exist between rodent and human cells (Fig. 4E).

## Discussion

The current study is directed at defining mechanisms that can contribute to regulating capacity for remyelination in the developed human CNS and that can potentially be therapeutically targeted. We have focused on the contribution of mature OLs and late progenitors (A2B5+O4+) present in human CNS. Such cells would also be relevant for myelin maintenance, a process that has a high turnover rate. The emphasis here is on the epigenomic state of these cells, using the active histone enhancer mark H3K27ac and chromatin accessibility to assess epigenomic access to sites relevant for myelination and for immune regulation. Our initial analyses focused on human OL lineage cells that we have isolated from surgically resected brain tissue samples from pediatric and adult age groups; for both mature OLs and late progenitors from these samples, we have shown that ensheathment capacity was significantly greater for the pediatric sample cells with both age and cell maturity contributing to the functional differences (17). Our previous studies showing differences between similar phenotypic cells derived from grey versus white matter from the same individuals support the contribution of maturity versus age alone (18). Our immunostaining data indicated that levels of the H3K27ac histone modification were higher in pediatric than adult samples, in both late progenitors and mature OLs. Levels were also decreased in the mature cell versus late progenitors in the adult derived samples. These results in human cells, contrast with observations in mice where no significant age-related differences in H3K27ac and H3K27me3 were detected between postnatal day 2 and postnatal day 60 (25).

To link function with epigenome, we first derived an overall signature of genes that were more expressed in the pediatric versus adult age group as ensheathment was superior for both late progenitor and mature cells for the younger age group. From this overall signature we could derive signatures that were relevant for OL differentiation and myelination (termed “myelination-related,” “immune-related”, and “stress/cell death processes”. All showed higher expression levels in the pediatric samples, compared to adult cells. Using data sets containing a full range of OL lineage cells (29–32), we could further show that those signatures showed an enrichment at earlier stages of oligodendrocyte lineage differentiation. Our analyses further revealed lower chromatin accessibility and H3K27ac levels in adult versus pediatric samples and in OL versus early lineage cells. The observation that the inhibitory histone mark H3K27me3 showed enrichment in the “myelination-related” signature in mature OLs versus OPCs, is consistent with observations that decreased progenitor marker expression is epigenetically regulated during OL differentiation.

Our analysis suggests the existence of species-specific differences in transcriptomic regulation between human and rodent oligodendrocytes, with rodent OPCs more effectively expressing transcriptional signatures related to myelination, immune response, and stress/cell death processes. In addition, our comparative analysis between OPC and OL in human and rodent cells, revealed significantly lower chromatin accessibility and H3K27ac in human mature OLs but only a trend for rodent cells, for all the transcriptional signatures analyzed. The levels of the repressive mark H3K27me3 were also lower for immune and age-related signatures, while they were significantly higher in OL for genes related to progenitor markers, included in the “myelination-related” transcriptional signature. Finally, while these species-specific epigenomic and transcriptomic changes are of interest, additional experiments would be necessary to determine whether and how each of the individual pathways contribute, positively or negatively, to differences in myelination potential and in response to immune and injury conditions between humans and rodents.

As part of the current study, we considered what may be the consequences of epigenomic modulation on susceptibility of OLs to injury especially in relation to interaction with the immune system as occurs in multiple sclerosis. We have shown that OL susceptibility to injury links both to age and maturity. Here we show that epigenome access is more apparent in the pediatric derived cells and raise the issue of whether modulation would have a similar effect. We submit the same applies to stress gene expression.

Overall, our data indicate the distinct myelination relevant epigenomic features of differentiated human oligodendrocyte lineage cells. The report of Liu et al using mature rodent OLs and in vivo demyelination models demonstrated that epigenome modulation alone (using deacetylase inhibitor ESI1) may provide an approach to enhance remyelination (24). We speculate whether in humans, such intervention alone would promote spontaneous remyelination or would be further enhanced by use of myelin promoting agents that would have better access to signaling pathways. Development of means to enhance myelination by modulating the epigenome will also need consider the coincident impact on immune reactivity.

## Materials and Methods

### Bulk RNA Sequencing Analysis and Signature Generation

RNA extracted from selected late progenitors (A2B5+) and mature OL (A2B5−) cells were subjected to bulk RNA sequencing at the Génome Québec Centre using an Illumina NovaSeq 6000 PE100 platform (Table S1). Raw reads underwent quality control, alignment, and quantification using the GenPipes workflow based on the GRCh38 human genome. Differential gene expression analysis was conducted with DESeq2 in R, employing normalization, variance stabilization, and transformation of raw counts (37). Differentially expressed genes (DEGs) were identified using thresholds of P<0.05 and log2 fold change >1.

To create the age-related signature, we overlapped differentially expressed genes from two comparisons (pediatric versus adult samples for both mature oligodendrocytes and late progenitor cells), identifying 1186 common genes. Non-coding genes were removed using Biomart, resulting in a final signature comprising 987 protein-coding genes. Using the same approach, we identified an age-upregulated signature containing 284 protein-coding genes.

Gene ontology (GO) analysis of significantly upregulated genes was performed using gProfiler, considering pathways with adjusted P<0.05 as significant (38). Results were visualized as bubble plots depicting the top GO biological process terms based on negative log10-transformed *P-values* and gene counts. Summary visualizations of GO biological processes were generated using Revigo with default parameters and represented in two-dimensional scatterplots (39).

Sub-signatures derived from the main age signature were generated based on GO results from gProfiler. We categorized significantly enriched terms into three main groups: oligodendrocyte lineage differentiation and myelination relevant (termed myelination related signature(MRS)), immune function (immune signature), and stress/cell death pathways (stress/cell death signature). Genes specific to each category were identified, removing overlaps between categories to ensure category-specific gene lists. These sub-signatures were analyzed alongside the primary age signature in subsequent analyses.

### Single nucleus/cell RNA-seq analysis

We analyzed three independent single-nucleus RNA-seq (snRNA-seq) datasets from Jakel et al. (2019), Absinta et al. (2021), and Trobisch et al. (2022) (29–31). For Trobisch et al. (2022), we utilized the authors’ provided data object. For the other studies, analysis was performed using the Seurat package (40). Gene expression data were imported via Read10X, creating Seurat objects requiring a minimum of three cells per gene and 200 features per cell. Cells were filtered based on nFeature\_RNA (200-2500) and mitochondrial content (percent.mt <20). Data were normalized (NormalizeData), variable features were identified (FindVariableFeatures, method “vst”, top 2000), and data scaling was applied (ScaleData). Dimensional reduction used RunPCA, neighbor graphs were created (FindNeighbors, dims=1:20), and clusters defined (FindClusters, resolution=0.1). UMAP visualization was performed (RunUMAP, dims=1:20). In addition to use original publication for each dataset to annotate clusters, we manually annotated and check clusters using known OL lineage markers (OPC markers: PTPRZ1, CSPG4, PDGFRA, SOX5, SOX6, GPR17; mature OL markers: MAG, MBP, MOG, PLP1, MOBP, CNP, MAL). Additionally, our previously published single-cell RNA-seq (scRNA-seq) dataset from cortical/subcortical regions, subventricular zones, and spinal cord samples was included, utilizing original normalized expression values (32).

### ChIP-seq, single-nuclei ATAC-seq, and nanoCUT&Tag analyses

ChIP-seq data for H3K27ac from Kozlenkov et al. (2024) (33), profiling human OPCs and mature oligodendrocytes from control postmortem samples, were analyzed. Normalized bigWig files (read counts per million) were visualized using Integrative Genomics Viewer (IGV) (41). IGV visualizations employed group autoscaling for comparable gene-specific views. H3K27ac was quantified within ±10 kb of transcription start sites (TSS) using VisR (RPM normalized) (42). These quantifications supported subsequent analyses including heatmaps and quartile calculations.

Single-nuclei ATAC-seq and nanoCUT&Tag datasets (H3K27ac and H3K27me3) from human and mouse samples (mouse snCUT&Tag) were analyzed similarly (34, 36). Data were loaded into VisR via IGV, quantified (RPKM) within ±10 kb of TSS regions for all genes, exported as matrices, and subsequently used for single-sample gene set enrichment analysis (ssGSEA) and visualization.

### Single-Sample Gene Set Enrichment Analysis (ssGSEA)

For ssGSEA, normalized gene expression data and gene signature files (GMT format) were loaded into pandas (43). Duplicate genes were averaged, and ssGSEA was executed using gseapy with rank normalization, zero permutations, and gene set sizes ranging from 15 to 15,000 (44). Enrichment scores were visualized using Prism, with significance determined by t-tests. Epigenomic ssGSEA analyses utilized similarly quantified accessibility (scATAC-seq), H3K27ac (ChIP-seq, nanoCUT&Tag), and H3K27me3 (nanoCUT&Tag) within ±10 kb of TSS (RPKM normalized). Human gene signatures were translated to mouse and rat orthologs for comparable analyses across species. For sn/scRNA-seq datasets, average expression levels of OPCs and mature oligodendrocytes were used for ssGSEA, pooling control samples within each study.

### Gene expression and H3K27ac clustering and heatmap visualization

Gene expression (normalized read counts) and H3K27ac (RPM) data were log2-transformed (numpy, adding 1 to avoid log-zero) and normalized (row-wise min-max scaling) (45). Custom diverging colormaps were generated with matplotlib (46). Hierarchical clustering (average linkage, correlation distance) and heatmap visualization were performed using seaborn’s clustermap, clustering either genes and samples or genes alone for specific analyses (47).

### Immunofluorescence based analysis of H3K27ac and H3K27me3 levels in human late progenitors (A2B5+) and mature OLs (A2B5−) in relation to age

For these studies, enriched populations of mature OLs (A2B5−) and late progenitor (A2B5+) cells were isolated from adult and pediatric surgically resected CUSA bag tissue samples as previously described (Luo et al 2022) using a trypsin and DNase digestion followed by Percoll gradient centrifugation procedure(17). The initial total dissociated cell sample was cultured in DMEM/F12 medium (Sigma, Oakville, ON, Canada) containing N1 (Sigma, Oakville, ON, Canada), 0.01% bovine serum albumin, and 1% penicillin–streptomycin (Invitrogen, Burlington, ON, Canada) overnight and the floating cells were collected, leaving behind the adherent microglia. A2B5 antibody conjugated microbeads (Miltenyi Biotec, Auburn, CA) were then used to select A2B5+ cells. Non-selected cells, referred to as A2B5− (or A−), comprise mature OLs. The late progenitors or A2B5+ (or A+) cells comprise ∼5-10% of the total OL fraction; the high majority (∼90%) express O4 (48, 49). As previously documented, these cell preparations are devoid of astrocytes and neurons and have a small proportion of microglia as documented by FACS (46,47) and single cell mRNA sequencing (Supp Fig 1 in Luo et al) (17). The 2 cells fractions (A+ and A-) express comparable levels of mature myelin genes (MBP and PLP) based on bulk RNA sequencing whereas progenitor genes (PDGFRa and PTPRZ1) are more highly expressed in the A+ cells (Fig 3 in Luo et al (16)). A2B5-positive and A2B5-negative cells were plated into 16-well poly-L-lysine- and extracellular matrix-coated chamber slides (30 000 cells per well) in defined medium containing DMEM/F12+ N1 medium supplemented with B27 (Life Technologies), PDGF-AA and bFGF. After 1-2 days in culture, cells were fixed and immunostained with Olig2 (Millipore, followed with goat-anti mouse Alexa Fluro-555 secondary antibody from Invitrogen) and H3K27ac (Abcam)/H3K27me3 (Diagenode) (followed by goat-anti rabbit Alexa Fluro-488 secondary antibody from Invitrogen) antibodies and DAPI as a nuclear counterstain. Intensity of H3K27ac/H3K27me3 nuclear immunoreactivity in Olig2+ cells was determined using confocal microscopy captured images (Table S2).

### Data, Materials, and Software Availability

We have used the following publicly available datasets from other research groups, including single nuclear RNA-seq datasets from Jakel et al. (2019, accession number: GSE118257), Absinta et al. (2021, accession number: GSE180759), and Trobisch et al. (2022, accession: https://cells.ucsc.edu/?ds=ms-cross-regional). We have used bulk RNA-seq data from young and aged rat from Neummann et al. (2019, accession number: GSE134765). For epigenomic datasets, we have used ChIP-seq dataset for H3K27ac from Kozlenkov et al. (2024, accession number: GSE239663), ATAC-seq modality of single cell multiomic of Velmeshev et al. (2023, accession: https://cells.ucsc.edu/?ds=pre-postnatal-cortex), human cell nanoCUT&Tag for H3K27ac and H3K27me3 as well single cell ATAC-seq data used from Kabbe et al. (2024, accession: https://cells.ucsc.edu/?ds=cns-nanocuttag-atac), Mouse Single-cell CUT&Tag for H3K27ac and H3K27me3 used from Bartosovic et al. (2021, accession: https://cells.ucsc.edu/?ds=mouse-brain-cutandtag), and Mouse adult cortex single cell ATAC-seq data originate from 10X dataset (https://support.10xgenomics.com/single-cell-atac/datasets/1.2.0/atac_v1_adult_brain_fresh_5k).

## Author contributions

AM. DD, QLC, PC, and JPA designed research; AM, DD, QLC and MY performed research; RD, JAS, SZ, TEK, MS, PC, and JPA contributed new reagents/analytic tools; AM, MY, QLC and DD analyzed data; and AM, JPA and PC wrote the paper.

## Supplementary Figures

**Figure S1.**
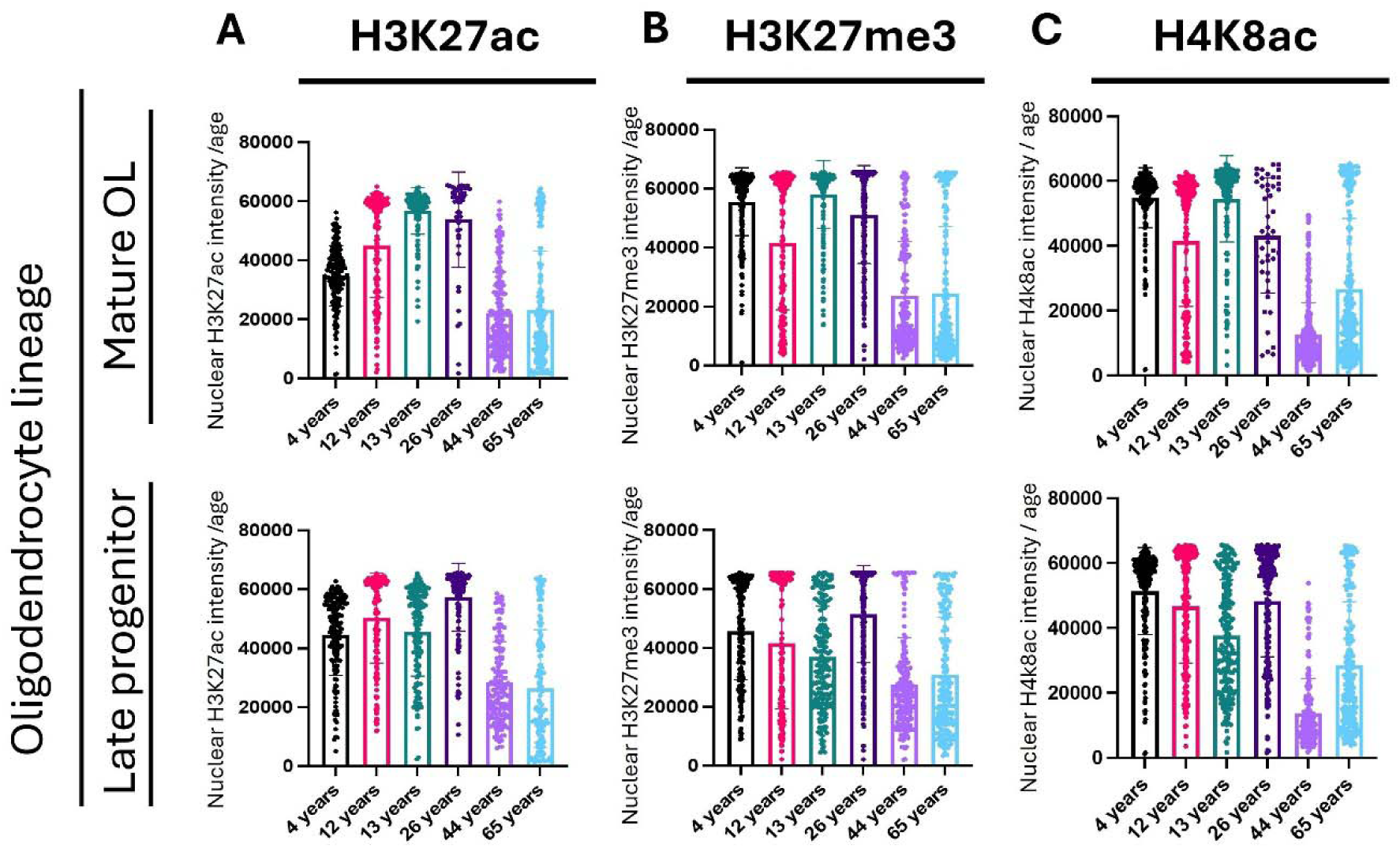
Adult mature oligodendrocytes (A2B5−) and late progenitor (A2B5+) cells have lower H3K27ac, H3K27me3, and H4K8ac levels compared with pediatric cells. **(A)** Bar graph of the nuclear intensity of H3K27ac immunoreactivity in OLIG2 cells over age. Data represent the H3K27ac immunoreactivity measured in a total of 900 cells from six donors. **(B)** Bar graph of the nuclear intensity of H3K27me3 immunoreactivity in OLIG2 cells over age. Data represent the H3K27me3 immunoreactivity measured in a total of 1200 cells from six donors. **(C)** Bar graph of the nuclear intensity of H4K8ac immunoreactivity in OLIG2 cells over age. Data represent the H4K8ac immunoreactivity measured in a total of 1200 cells from six donors.

**Figure S2.**
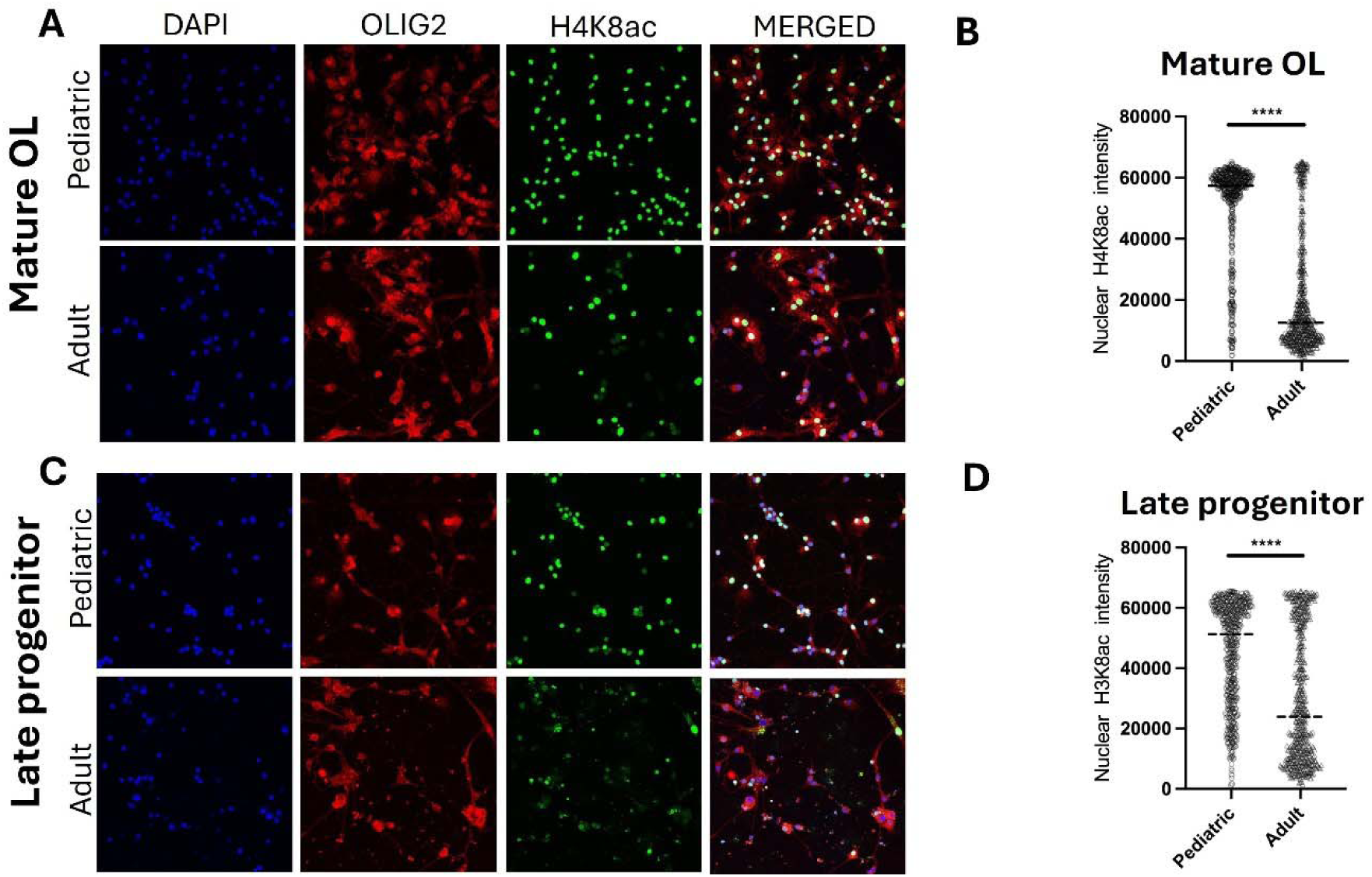
Changes in H4K8ac nuclear intensity of A2B5− and A2B5+ cells over age. **(A)** Confocal image of A2B5− cells obtained from pediatric and adult human patients, stained with OLIG2 antibodies (red), H4K8ac antibodies (green) and DAPI (blue) as a nuclear counterstain. **(B)** Violin plots of the nuclear intensity of H3K27me3 immunoreactivity in OLIG2^+^ cells in pediatric and adult brains. Data represent the H4K8ac immunoreactivity measured in a total of 600 cells from three biological replicates (****P < 0.0001, unpaired t-test). **(C)** Confocal image of A2B5+ cells obtained from pediatric and adult human patients, stained with OLIG2 antibodies (red), H4K8ac antibodies (green) and DAPI (blue) as a nuclear counterstain. **(D)** Violin plots of the nuclear intensity of H4K8ac immunoreactivity in OLIG2^+^ cells in pediatric and adult brains. Data represent the H4K8ac immunoreactivity measured in a total of 600 cells from three biological replicates (****P < 0.0001, unpaired t-test).

**Figure S3.**
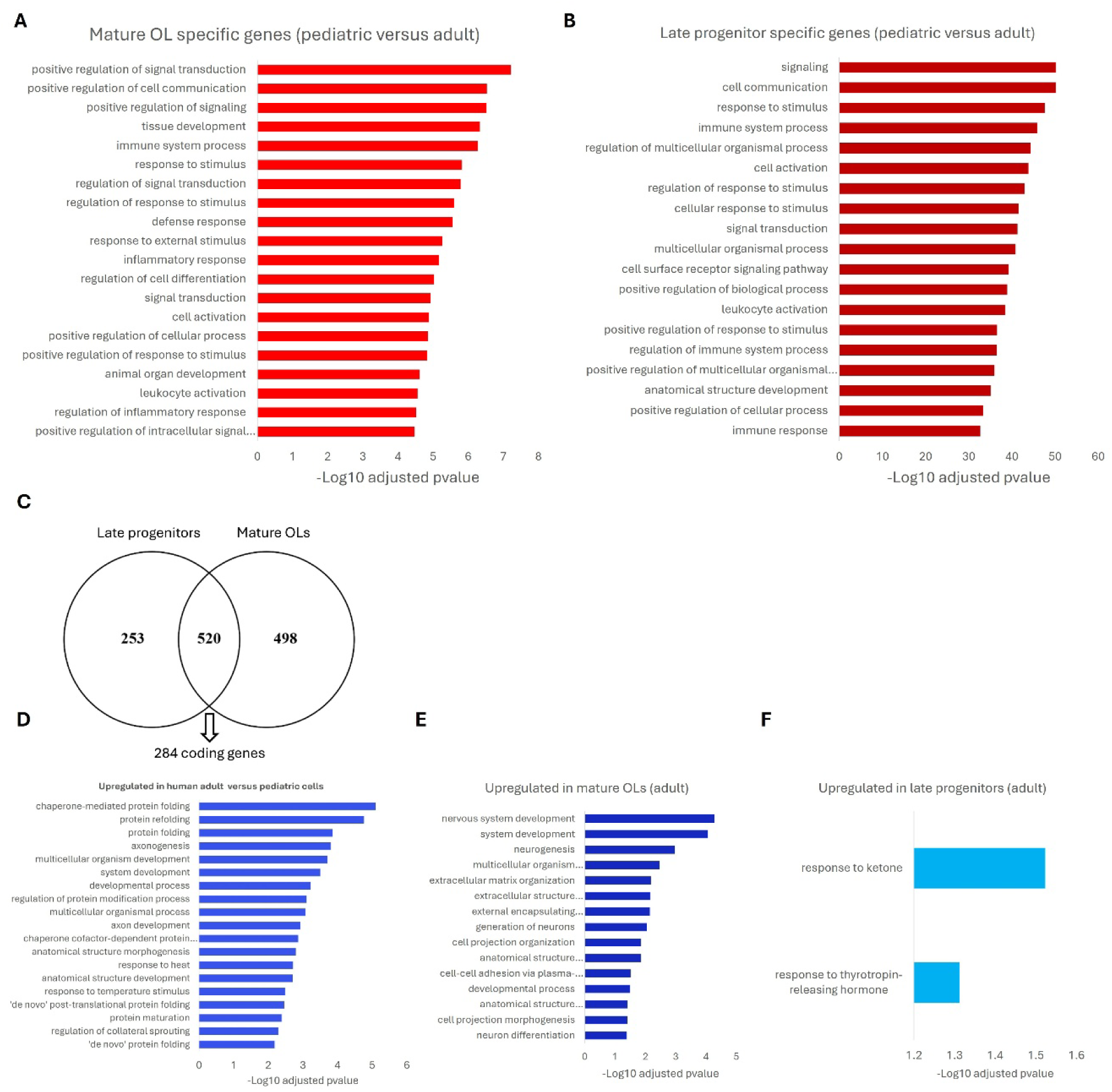
Common and specific transcriptomic profiles downregulated and upregulated in mature oligodendrocytes and late progenitors in adult samples compared to pediatric samples. **(A)** Gene ontology using gProfiler on the genes upregulated (log2FC > 1 and p < 0.05) in mature OLs and not in late progenitors in pediatric samples versus adult samples. Top 20 biological processes were selected based on the adjusted p-values and visualized. **(B)** Gene ontology using gProfiler on the genes upregulated (log2FC > 1 and p < 0.05) in the late progenitors OLs and not in mature OLs in pediatric samples versus adult samples. Top 20 biological processes were selected based on the adjusted p-values and visualized. **(C)** Age-related transcriptomic signatures were identified by comparing adult (>18 years) and pediatric (<18 years) samples in two distinct human oligodendrocyte lineage populations: mature oligodendrocytes (A2B5−) and late oligodendrocyte progenitors (A2B5+). Genes upregulated in adult samples for each comparison were filtered based on statistical significance (p < 0.05) and log2 fold-change (>1), resulting in a set of differentially expressed genes. Intersection of these sets yielded 520 overlapping genes; removal of non-coding genes refined this list to 284 protein-coding genes. **(D)** Gene ontology analysis was performed using gProfiler on the 284 genes, and top significant GO terms (adjusted p < 0.05) were visualized. (**E)** Gene ontology using gProfiler on the downregulated genes (log2FC > 1 and p < 0.05) in mature OLs and not in late progenitors in pediatric samples versus adult samples. All biological processes were selected based on the adjusted p-values and visualized. **(F)** Gene ontology using gProfiler on the genes downregulated (log2FC > 1 and p < 0.05) in late progenitors and not in mature cells in pediatric samples versus adult samples. All biological processes were selected based on the adjusted p-values and visualized.

**Figure S4.**
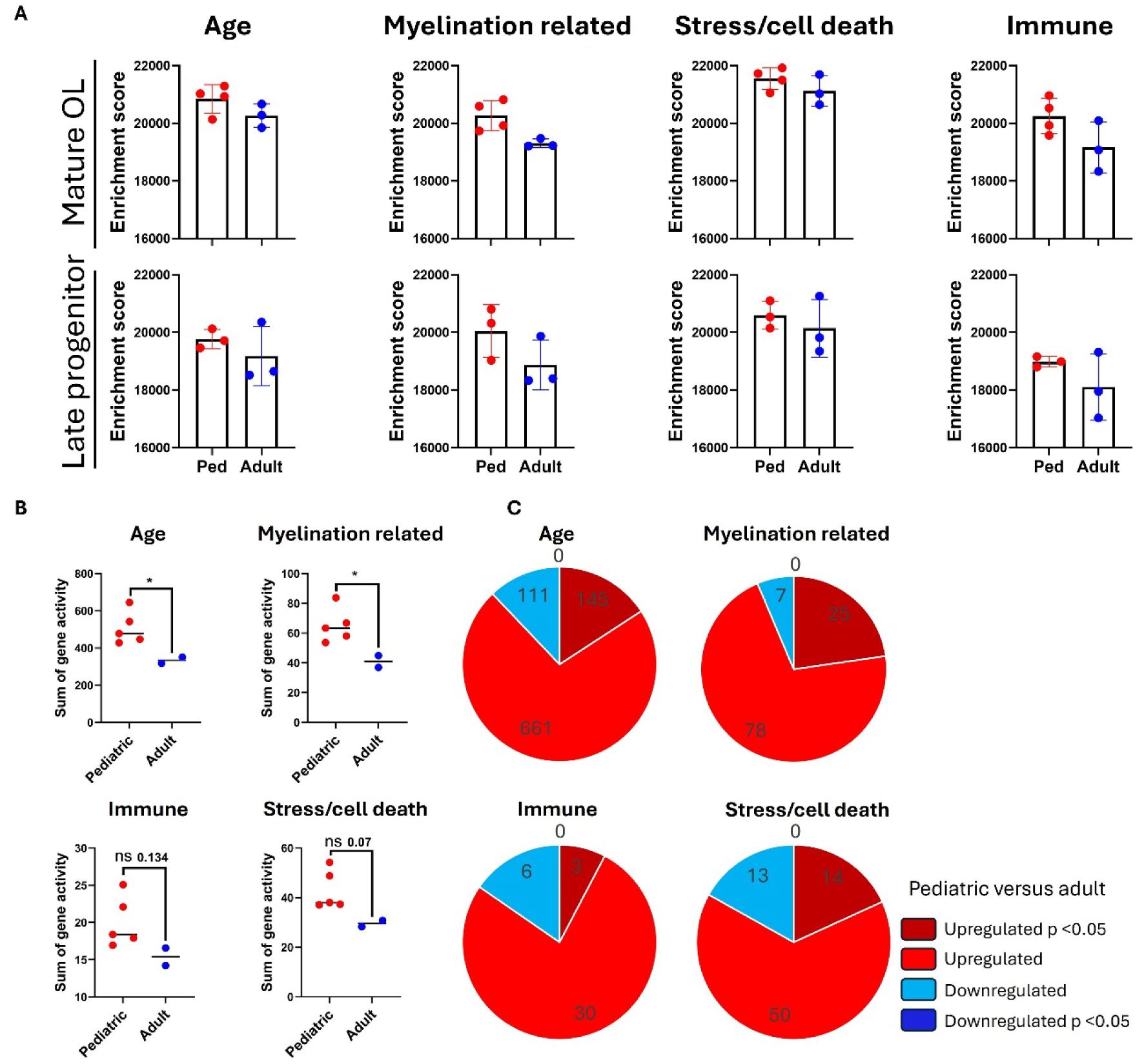
Transcriptomic signature downregulated in oligodendrocyte lineage cells after one day in culture shows a similar trend at six days. **(A)** Normalized read counts from RNA-seq data of day 6 cells in culture were used (Mohammadnia et al., 2024). ssGSEA was conducted on mature oligodendrocyte and late progenitor expression profiles for the identified gene categories (full age-related signature, myelination related, immune, and stress/cell death signatures). Enrichment scores were plotted, and statistical significance between pediatric and adult groups was assessed using t-tests. All signatures showed significantly higher enrichment scores in pediatric samples compared to adults. **(B)** The ATAC-seq modality of the single-cell multiomic dataset for OPCs from Velmeshev et al. (2023), including pediatric (ages 1–8 years old) and adult (25 and 39 years old) donors, was used to assess accessibility (gene activity) of gene signatures derived from comparing bulk RNA-seq adult human primary cells with pediatric counterparts in OPCs. For each sample, average gene activity for genes in each group was calculated, and the sum of gene activities per signature per sample was used. A two-sided t-test was used to assess the significance of differences between adults and pediatrics; ** and *** indicate p < 0.005 and p < 0.0005, respectively. **(C)** We performed a pseudobulk differential analysis on OPCs by aggregating normalized gene-activity values (log(CP10k)) to donor-level means, using Individual as the biological replicate. Donors were split into Early and Adult groups, and per-gene differences were tested with Welch’s t-test to accommodate unequal variances and sample sizes. P values were corrected for multiple testing using Benjamini–Hochberg FDR, yielding differentially active genes per cell type. Genes from four signatures were extracted and divided into four categories: significantly upregulated (p < 0.05, dark red), upregulated (non-significant, light red), downregulated (non-significant, light blue), and significantly downregulated (p < 0.05, dark blue).

**Figure S5.**
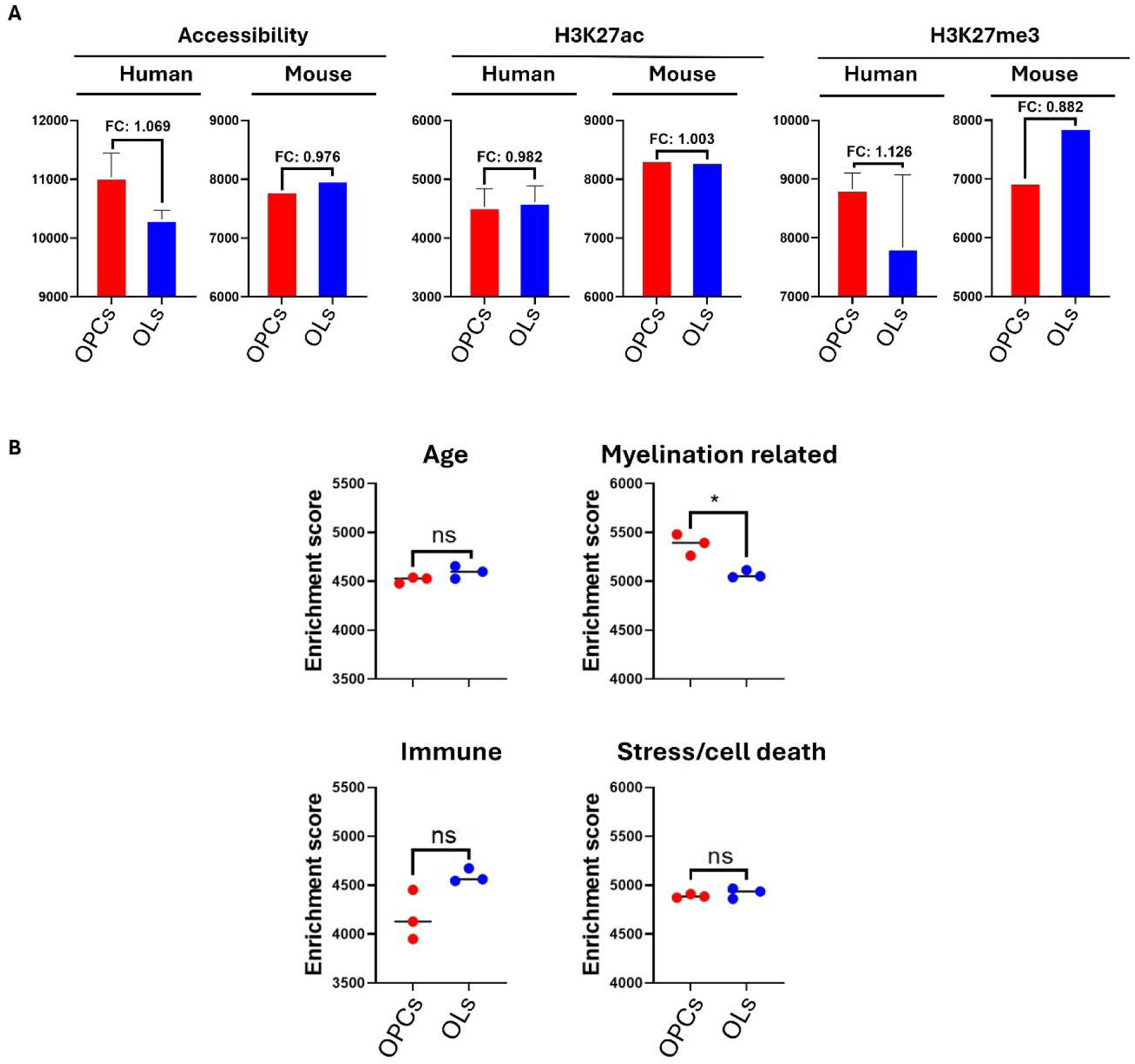
Comparative epigenomic and transcriptome profiling in human and rodents. **(A)** Single-cell ATAC-seq and single-cell nanoCUT&Tag datasets for human (Kabbe et al., 2024) and single-cell ATAC-seq and snCUT&Tag datasets for mouse (Bartosovic et al., 2021) were used. Accessibility, H3K27ac (activation mark), and H3K27me3 (repressive mark) were quantified within ±10 kb TSS regions across all genes (normalized as RPKM values) in OPCs and mature oligodendrocytes from both species. ssGSEA was performed for stress/cell death gene signature. Enrichment scores are plotted with fold differences plotted for the conditions. Red represents OPCs and blue indicates mature cells. **(B)** SsGSEA of rat transcriptomes comparing young (2–3 months old) and aged (22–24 months old) samples was performed using normalized read counts generated from raw data obtained from Neumann et al. (2019). Among the four signatures analyzed (age, myelination related, immune, and stress/cell death), only the myelination related signature showed significant enrichment in young versus aged samples (unpaired t-test). Red and blue denote young and aged groups, respectively.

**Table S1.** The Table summarizes the age, sex, and diagnosis of the samples used for bulk RNA-seq dataset.

|  | **ID** | **Age** | **Sex** | **Diagnosis** |
| --- | --- | --- | --- | --- |
| **Pediatric** | HA811 | 2 years | M | Focal cortical dysplasia |
|  | HA785 | 4 years | M | Ganglioma |
|  | HA810 | 8 years | M | Focal cortical dysplasia |
| **Adult** | HA777 | 26 years | F | Gliosis |
|  | HA789 | 21 years | M | Focal cortical dysplasia |
|  | HA791 | 38 years | M | Gliosis |

**Table S2.**
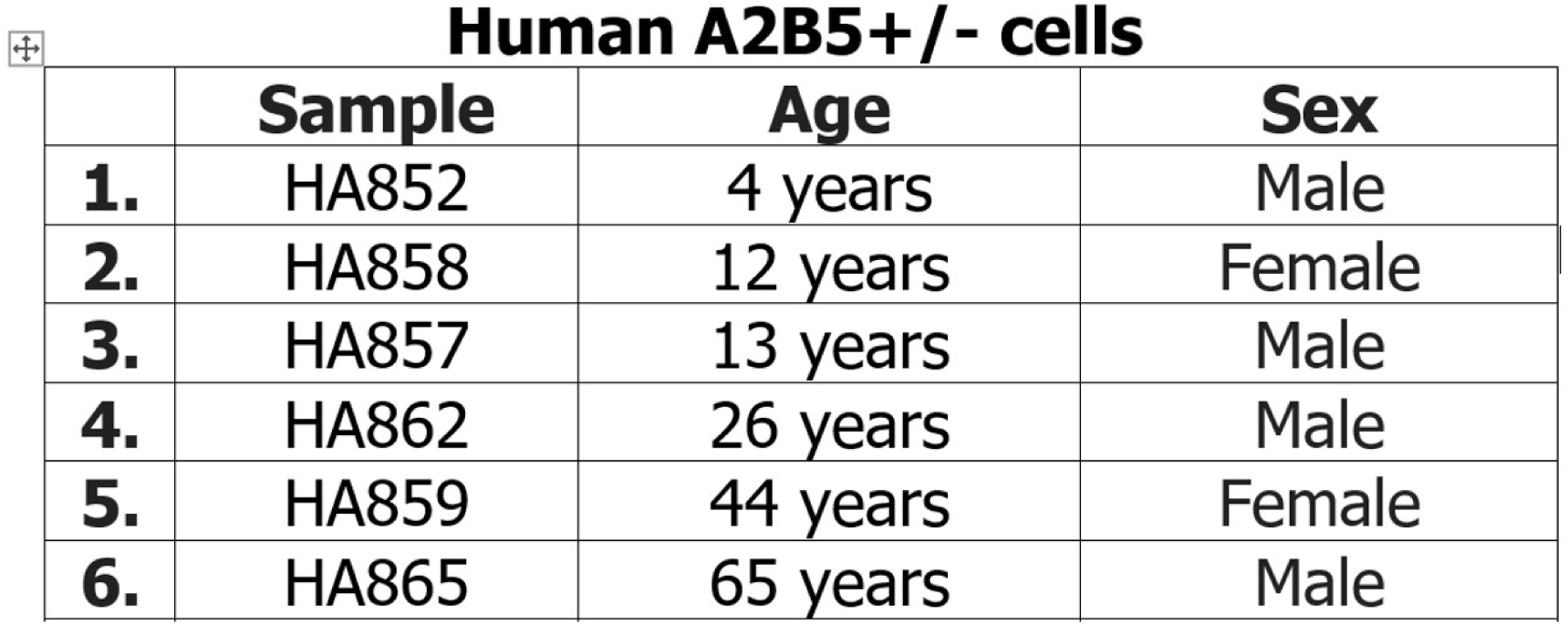
The Table summarizes the age, sex, and diagnosis of the samples used for immunofluorescence staining.

**Human A2B5+/- cells**
|  | **Sample** | **Age** | **Sex** |
| --- | --- | --- | --- |
| **1.** | HA852 | 4 years | Male |
| **2.** | HA858 | 12 years | Female |
| **3.** | HA857 | 13 years | Male |
| **4.** | HA862 | 26 years | Male |
| **5.** | HA859 | 44 years | Female |
| **6.** | HA865 | 65 years | Male |

## References

1. M. Simons, K. A. Nave, Oligodendrocytes: Myelination and axonal support. Cold Spring Harb. Perspect. Biol. 8 (2016).

2. K. Heß, et al., Lesion stage-dependent causes for impaired remyelination in MS. Acta Neuropathol. 140, 359–375 (2020).

3. M. Albert, J. Antel, W. Brück, C. Stadelmann, Extensive cortical remyelination in patients with chronic multiple sclerosis. Brain Pathol. 17, 129–138 (2007).

4. J. W. L. Brown, et al., Remyelination varies between and within lesions in multiple sclerosis following bexarotene. Ann. Clin. Transl. Neurol. 9, 1626–1642 (2022).

5. A. Chang, et al., Cortical remyelination: A new target for repair therapies in multiple sclerosis. Ann. Neurol. 72, 918–926 (2012).

6. E. M. M. Strijbis, E. J. Kooi, P. van der Valk, J. J. G. Geurts, Cortical remyelination is heterogeneous in multiple sclerosis. J. Neuropathol. Exp. Neurol. 76, 390–401 (2017).

7. A. Lazzarotto, et al., Time is myelin: early cortical myelin repair prevents atrophy and clinical progression in multiple sclerosis. Brain 147, 1331–1343 (2024).

8. S. A. G. Robin J.M. Franklin, Benedetta Bodini, Remyelination in the Central Nervous System. Cold Spring Harb. Perspect. Biol. 7, 1–28 (2015).

9. M. S. Y. Yeung, et al., Dynamics of oligodendrocyte generation and myelination in the human brain. Cell 159, 766–774 (2014).

10. I. D. Duncan, et al., The adult oligodendrocyte can participate in remyelination. Proc. Natl. Acad. Sci. U. S. A. 115, E11807–E11816 (2018).

11. A. Mezydlo, et al., Remyelination by surviving oligodendrocytes is inefficient in the inflamed mammalian cortex. Neuron 111, 1748–1759.e8 (2023).

12. R. C. Armstrong, H. H. Dorn, C. V. Kufta, E. Friedman, M. E. Dubois-Dalcq, Pre-oligodendrocytes from adult human CNS. J. Neurosci. 12, 1538–1547 (1992).

13. M. S. Windrem, et al., Fetal and adult human oligodendrocyte progenitor cell isolates myelinate the congenitally dysmyelinated brain. Nat. Med. 10, 93–97 (2004).

14. N. S. Roy, et al., Identification, isolation, and promoter-defined separation of mitotic oligodendrocyte progenitor cells from the adult human subcortical white matter. J. Neurosci. 19, 9986–9995 (1999).

15. L. Kirby, et al., Oligodendrocyte precursor cells present antigen and are cytotoxic targets in inflammatory demyelination. Nat. Commun. 10, 1–20 (2019).

16. Q. L. Cui, et al., Oligodendrocyte progenitor cell susceptibility to injury in multiple sclerosis. Am. J. Pathol. 183, 516–525 (2013).

17. J. X. X. Luo, et al., Human Oligodendrocyte Myelination Potential; Relation to Age and Differentiation. Ann. Neurol. 91, 178–191 (2022).

18. Q. Cui, et al., Myelination potential and injury susceptibility of grey versus white matter human oligodendrocytes. Brain 148, 921–932 (2025).

19. B. Emery, T. L. Wood, Regulators of Oligodendrocyte Differentiation. Cold Spring Harb. Perspect. Biol. 16 (2024).

20. S. Copray, J. L. Huynh, F. Sher, P. Casaccia-Bonnefil, E. Boddeke, Epigenetic mechanisms facilitating oligodendrocyte development, maturation, and aging. Glia 57, 1579–1587 (2009).

21. J. Liu, et al., Chromatin landscape defined by repressive histone methylation during oligodendrocyte differentiation. J. Neurosci. 35, 352–365 (2015).

22. S. Shen, et al., Age-dependent epigenetic control of differentiation inhibitors is critical for remyelination efficiency. Nat. Neurosci. 11, 1024–1034 (2008).

23. X. Pedre, et al., Changed histone acetylation patterns in normal-appearing white matter and early multiple sclerosis lesions. J. Neurosci. 31, 3435–3445 (2011).

24. X. Liu, et al., Small-molecule-induced epigenetic rejuvenation promotes SREBP condensation and overcomes barriers to CNS myelin regeneration. Cell 187, 2465–2484.e22 (2024).

25. D. K. Dansu, et al., Histone H4 acetylation differentially modulates proliferation in adult oligodendrocyte progenitors. J. Cell Biol. 223 (2024).

26. A. Mohammadnia, et al., Age-dependent effects of metformin on human oligodendrocyte lineage cell ensheathment capacity. Brain Commun. 6, 1–13 (2024).

27. D. Velmeshev, et al., Single-cell analysis of prenatal and postnatal human cortical development. Science (80-.). 382 (2023).

28. C. A. Herring, et al., Human prefrontal cortex gene regulatory dynamics from gestation to adulthood at single-cell resolution. Cell 185, 4428–4447.e28 (2022).

29. S. Jäkel, et al., Altered human oligodendrocyte heterogeneity in multiple sclerosis. Nature 566, 543–547 (2019).

30. M. Absinta, et al., A lymphocyte–microglia–astrocyte axis in chronic active multiple sclerosis. Nature 597, 709–714 (2021).

31. T. Trobisch, et al., Cross-regional homeostatic and reactive glial signatures in multiple sclerosis. Acta Neuropathol. 144, 987–1003 (2022).

32. M. Yaqubi, et al., Regional and age-related diversity of human mature oligodendrocytes. Glia 70, 1938–1949 (2022).

33. A. Kozlenkov, et al., Novel method of isolating nuclei of human oligodendrocyte precursor cells reveals substantial developmental changes in gene expression and H3K27ac histone modification. Glia 72, 69–89 (2024).

34. M. Kabbe, et al., Single-nuclei histone modification profiling of the adult human central nervous system unveils epigenetic memory of developmental programs. bioRxiv 2024.04.15.589512 (2024).

35. B. Neumann, et al., Metformin Restores CNS Remyelination Capacity by Rejuvenating Aged Stem Cells. Cell Stem Cell 25, 473–485.e8 (2019).

36. M. Bartosovic, M. Kabbe, G. Castelo-Branco, Single-cell CUT&Tag profiles histone modifications and transcription factors in complex tissues (Springer US, 2021).

37. M. I. Love, W. Huber, S. Anders, Moderated estimation of fold change and dispersion for RNA-seq data with DESeq2. Genome Biol. 15, 1–21 (2014).

38. U. Raudvere, et al., G:Profiler: A web server for functional enrichment analysis and conversions of gene lists (2019 update). Nucleic Acids Res. 47, W191–W198 (2019).

39. F. Supek, M. Bošnjak, N. Škunca, T. Šmuc, Revigo summarizes and visualizes long lists of gene ontology terms. PLoS One 6 (2011).

40. A. Butler, P. Hoffman, P. Smibert, E. Papalexi, R. Satija, Integrating single-cell transcriptomic data across different conditions, technologies, and species. Nat. Biotechnol. 36, 411–420 (2018).

41. H. Thorvaldsdóttir, J. T. Robinson, J. P. Mesirov, Integrative Genomics Viewer (IGV): high-performance genomics data visualization and exploration. Brief. Bioinform. 14, 178–192 (2013).

42. H. Younesy, T. Möller, M. C. Lorincz, M. M. Karimi, S. J. M. Jones, VisRseq: R-based visual framework for analysis of sequencing data. BMC Bioinformatics 16, 1–14 (2015).

43. W. McKinney, Data Structures for Statistical Computing in Python. Proc. 9th Python Sci. Conf. 1, 56–61 (2010).

44. Z. Fang, X. Liu, G. Peltz, GSEApy: a comprehensive package for performing gene set enrichment analysis in Python. Bioinformatics 39, 1–3 (2023).

45. C. R. Harris, et al., Array programming with NumPy. Nature 585, 357–362 (2020).

46. J. D. Hunter,: a 2D G. Comput. Sci. Eng. 9, 90–95 (2007).

47. M. Waskom, Seaborn: Statistical Data Visualization. J. Open Source Softw. 6, 3021 (2021).

48. C. Esmonde-White, et al., Distinct function-related molecular profile of adult human A2B5-positive pre-oligodendrocytes versus mature oligodendrocytes. J. Neuropathol. Exp. Neurol. 78, 468–479 (2019).

49. S. Y. Leong, et al., Heterogeneity of oligodendrocyte progenitor cells in adult human brain. Ann. Clin. Transl. Neurol. 1, 272–283 (2014).

